# Inhibition of mitochondrial permeability transition by deletion of the ANT family and CypD

**DOI:** 10.1101/506964

**Authors:** Jason Karch, Michael J. Bround, Hadi Khalil, Michelle A. Sargent, Nadina Latchman, Naohiro Terada, Pablo M. Peixoto, Jeffery D. Molkentin

## Abstract

The mitochondrial permeability transition pore (MPTP) has resisted molecular identification for decades. The original model of the MPTP had the adenine nucleotide translocator (ANT) as the inner membrane pore-forming component. Indeed, reconstitution experiments showed that recombinant or purified ANT generates MPTP-like pores in lipid bilayers. This model was challenged when mitochondria from *Ant1/2* double null mouse liver still showed MPTP activity. Because mice contain and express 3 *Ant* genes, here we reinvestigated the genetic basis for the ANTs as comprising the MPTP. Liver mitochondria from *Ant1, Ant2*, and *Ant4* deficient mice were highly refractory to Ca^2+^-induced MPT, and when also given cyclosporine A, MPT was completely inhibited. Moreover, liver mitochondria from mice with quadruple deletion of *Ant1, Ant2, Ant4* and *Ppif* (cyclophilin D, target of CsA) lacked Ca^2+^-induced MPT. Finally, inner membrane patch clamping in mitochondria from *Ant1, Ant2* and *Ant4* triple null mouse embryonic fibroblasts (MEFs) showed a loss of MPT-like pores. Our findings suggest a new model of MPT consisting of two distinct molecular components, one of which is the ANTs and the other of which is unknown but requires CypD.

**One Sentence Summary:** Genetic deletion of *Ant1/2/4* and *Ppif* in mice fully inhibits the mitochondrial permeability transition pore

---

Mitochondrial permeability transition pore (MPTP) opening contributes to various pathologies involving necrotic cell death after ischemic injuries or degenerative muscle and brain diseases (*1–5*). The MPTP is a <1.5 kDa pore within the inner mitochondrial membrane which opens in the presence of high matrix Ca^2+^ and/or ROS (*6, 7*). Although numerous regulators of this pore have been identified, the molecular pore forming component of the MPTP remains undefined (*8, 9*). The adenine nucleotide translocator (ANT) family was originally proposed to be the pore forming component of the MPTP due to the fact that ANT ligands and inhibitors have profound effects on the opening of the MPTP and that reconstituted ANTs in lipid bilayers are able to form channels similar to that of the MPTP (*10–15*). However, this model was largely rejected when it was reported that mouse liver mitochondria lacking the genes Ant1 and Ant2, were able to undergo permeability transition, albeit this transition required substantially higher levels of Ca^2+^ to engage (*16*). Here we investigated whether the third Ant gene in mice, *Ant4*, compensates for the loss of *Ant1* and *Ant2* and whether mitochondria completely null for all ANT isoforms would be able to undergo permeability transition (humans contain 4 Ant genes (*Ant1-4*) while mice lack *Ant3*) (*17*). More recently a model in which the mitochondrial F_1_F_O_-ATPase serves at the MPTP has been proposed (*18–20*), although not without controversy as some data dispute the role of the F_1_F_0_-ATPase (*21–23*), thus leaving the MPTP undefined.

We first characterized the tissue distribution of all 3 ANT isoforms in the mouse by Western blotting from mitochondria lysates. In wild-type (WT) mice, ANT1 is the major isoform expressed in the heart and skeletal muscle, while ANT2 is expressed in all tissues examined besides the testis, which predominantly expresses ANT4 (Fig. 1A). Importantly, at baseline mouse liver only detectably expresses ANT2 (Fig. 1A). In mice globally lacking either Ant1 or Ant4, the ANT2 isoform becomes upregulated to compensate for their loss (Fig. 1A). Importantly, we observed that ANT4 expression is induced in liver mitochondria from mice lacking ANT1 and ANT2 (Fig. 1B). This observation of compensation by ANT4 when *Ant1* and *Ant2* were deleted from the liver likely explains why Kokoszka and colleagues failed to observe a more severe loss of MPTP activity (*16*). The *Ant2* gene was LoxP-targeted to permit tissue specific deletion because full somatic *Ant2*^-/-^ mice are embryonic lethal (*24*).

**Fig. 1.**
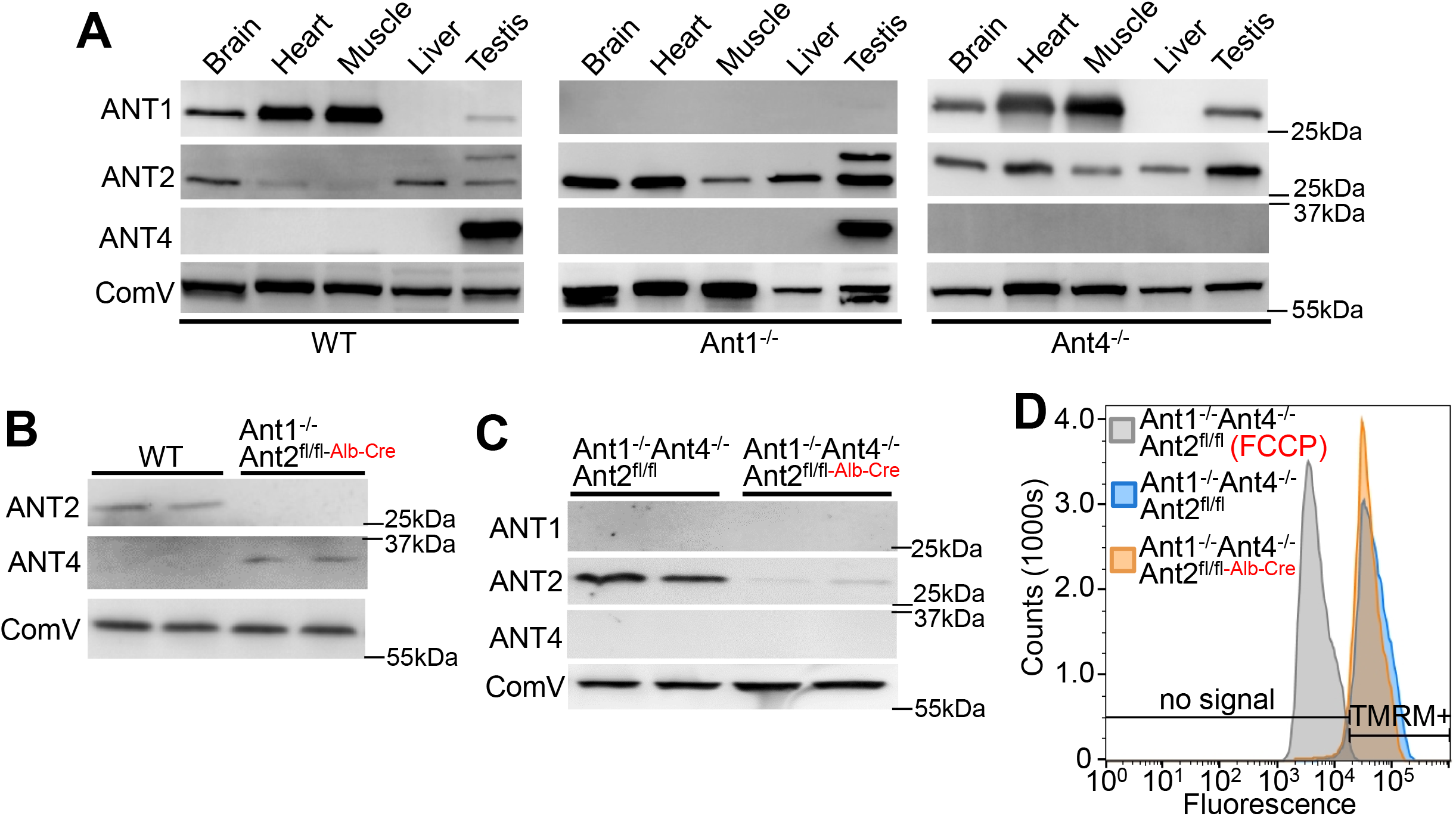
Tissue specific expression and compensation of the ANT family. (**A**) Western blots of isolated mitochondria from various mouse tissues for ANT1, ANT2, ANT4 and a loading control for complex V (ComV)). ANT2 is upregulated in all 5 tissues analyzed to compensate for the loss of the *Ant1* gene, or ANT2 is upregulated in testis in response to loss of the *Ant4* gene. These results are representative of 3 independent experiments. (**B**) Western blots for ANT2 and ANT4 from isolated liver mitochondria from the indicated gene-targeted mice, showing that ANT4 is induced for the first time in the absence of the *Ant1* and *Ant2* genes. A loading control for complex V is shown (ComV). These results are representative of 3 independent experiments. (**C**) Western blots of ANT1, ANT2, ANT4, and a loading control for complex V, performed using extracts of isolated mitochondria from *Ant1^-/-^ Ant4^-/-^ Ant2^fl/fl^* (only ANT2 is remaining) and *Ant1^-/-^ Ant4^-/-^ Ant2^fl/fl-Alb-Cre^* (*Ant* triple null livers). Data are representative of 3 independent experiments. (**D**) Representative FACS analysis of TMRE stained mouse liver mitochondria isolated from *Ant1^-/-^ Ant4^-/-^ Ant2^fl/fl^* versus *Ant1^-/-^ Ant4^-/-^ Ant2^fl/fl-Alb-Cre^* mice. The TMRE fluorescence threshold is shown in the bottom of the panel versus when no fluorescence signal is detected above background. FCCP is a potent uncoupler given to depolarize mitochondria. Results are representative of 3 independent experiments.

Here we generated mice null for all ANT isoforms in the liver by crossing total somatic *Ant1^-/-^, Ant4*^-/-^ with *Ant2-loxP* (fl) mice with an albumin promoter driven Cre transgenic mouse line (*Ant2^fl/fl-Alb-Cre^*) (*16, 25*). The loss of all ANT isoform expression in the liver was confirmed by Western blot analysis on isolated liver mitochondria (Fig. 1C). In all liver-based experiments *Ant1^-/-^ Ant4^-/-^ Ant2^fl/fl^* mice were used as controls because ANT2 is the overwhelming isoform expressed in liver (Fig. 1A and C). More importantly, this strategy allowed us to directly compare littermates (*Ant1^-/-^ Ant4^-/-^ Ant2^fl/fl-Alb-Cre^* versus *Ant1^-/-^ Ant4^-/-^ Ant2^fl/fl^*). TMRE staining showed comparable membrane potential that was sensitive to FCCP treatment in mitochondria from triple null livers and the ANT2 only expressing controls (*Ant1^-/-^ Ant4^-/-^ Ant2^fl/fl^*) (Fig. 1D). H&E-stained histological liver sections and transmission electron microscopy also showed no overt pathology in *Ant* triple nulls versus ANT2 only expressing mice (Fig. S1A). Surprisingly, *Ant* triple null mitochondria from liver also showed no difference in stimulated and maximum oxygen consumption rate compared with ANT2 only, suggesting ANT function is somehow compensated in these mitochondria (Fig. S1B).

To investigate the status of MPTP activity we used the 2 most commonly employed MPTP assays in isolated mitochondria: that of Ca^2+^ retention capacity (CRC) using step-wise 40 μM Ca^2+^ additions over time, and that of absorbance-based assessment of mitochondrial swelling. Mitochondria from WT mouse liver only took up 3 stepwise Ca^2+^ additions before opening and dumping their Ca^2+^ into the solution, which contains a fluorescence indictor (Fig. 2A). This Ca^2+^-induced MPTP opening caused immediate mitochondria swelling indicated by a decrease in light absorbance of these organelles in solution (Fig. 2B). Consistent with previous results (*16*), loss of ANT1 and ANT2 desensitized Ca^2+^-induced MPTP opening similarly to the level of WT mitochondria treated with cyclosporine A (CsA) (*26, 27*), which is an inhibitor of CypD that regulates the MPTP (Fig. 2A and B). Mitochondria lacking all isoforms of the ANT family are substantially further desensitized to Ca^2+^-induced MPTP opening, however, the MPTP can still open with extreme levels of Ca^2+^ (Fig. 2A and B), confirming that the ANT family is still dispensable for MPTP opening in liver. However, triple null mitochondria also treated with CsA showed Ca^2+^ uptake to the physical limits of the assay at 4.8 mM before mitochondria stopped taking up Ca^2+^, although mitochondria swelling never occurred (Fig. 2A and B), unless the non-specific permeabilizing agent alamethicin was added (Fig. 2B, arrowhead).

**Fig. 2.**
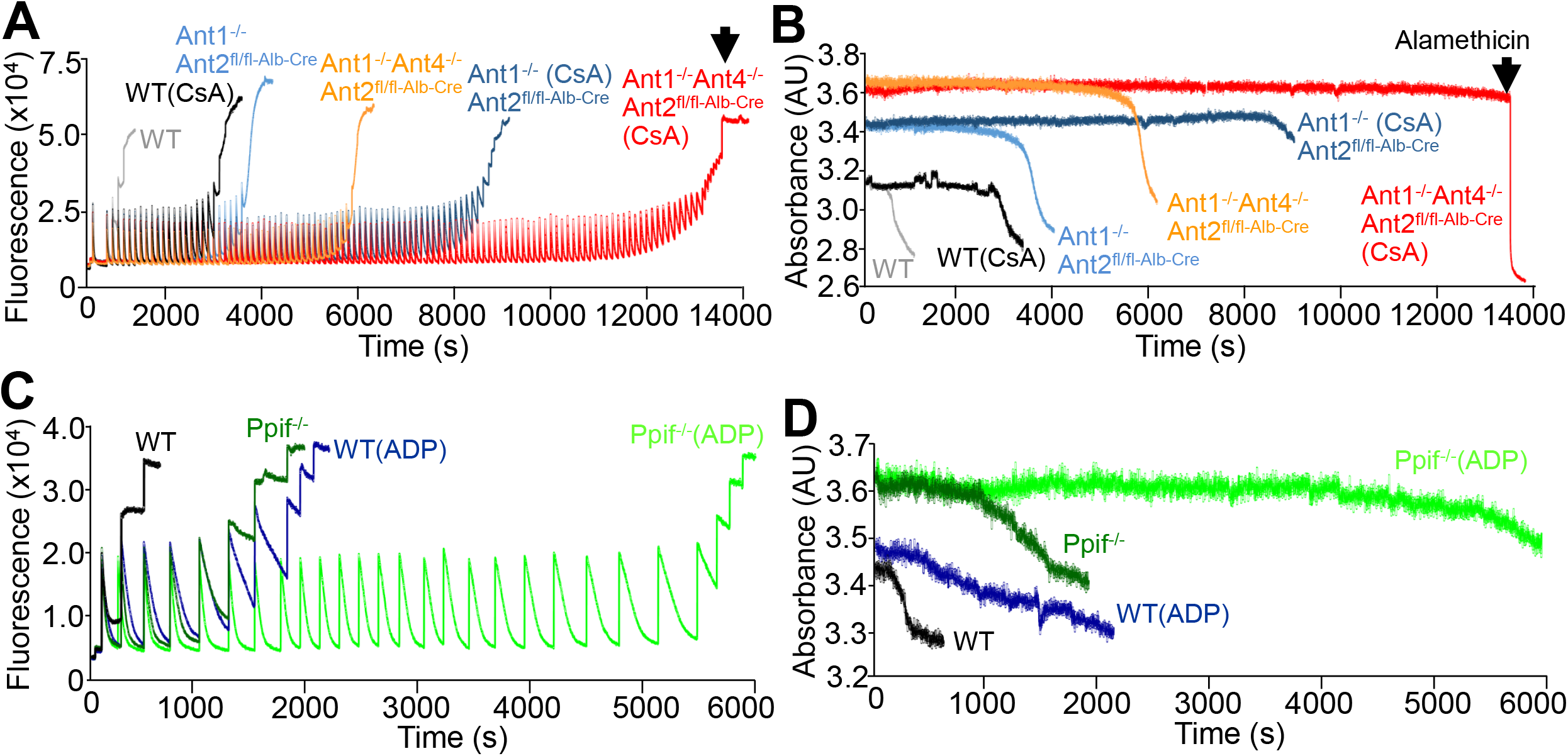
Loss of the Ant family or CypD desensitizes the MPTP, while Ant family deletion with CsA completely inhibits MPTP opening. (**A**) Ca^2+^ retention capacity assay with Calcium Green-5N fluorescence indicator in the buffer was performed in purified liver mitochondria isolated from wild-type (**WT**), *Ant1^-/-^ Ant2^fl/fl-Alb-Cre^*, or *Ant1^-/-^ Ant4^-/-^ Ant2^fl/fl-Alb-Cre^* mice treated with our without 5 μM cyclosporine A (CsA). Two milligrams of mitochondria were used in each assay and 40 μM pulses of CaCl_2_ were continually given (represented by each peak in fluorescence in the traces) until mitochondrial either underwent MPTP or simply had saturated Ca^2+^ uptake. The arrowhead shows alamethicin addition, a membrane permeabilizing agent. (**B**) Simultaneous with the assay shown in “A”, light absorbance was recorded during the course of Ca^2+^ additions to measure mitochondrial swelling represented by a decrease in absorbance. The results shown are representative traces from at least 3 independent assays. (**C**) Ca^2+^ retention capacity assay using similar conditions as in “A” performed on liver mitochondria isolated from WT or *Ppif*^-/-^ mice treated with or without 500 nM ADP. (**D**) Absorbance-based mitochondria swelling was simultaneously measured at the same time in the samples shown in “C”. The results shown are representative traces from at least 3 independent assays.

We observed a similar synergistic effect in the reciprocal experiment when *Ppif*^-/-^ mitochondria lacking CypD were treated with ADP, a known desensitizer of the MPTP and ligand for the ANTs (Fig. 2C and D). Together these data reveal that MPTP requires at least one ANT family member or CypD. These studies further suggest that there may be at least two separate channels which comprise the total MPTP activity in liver: one channel consisting of the ANT family that can escape CypD regulation and another that is absolutely dependent on CypD to open.

To further address the data obtained in purified mitochondria with CsA in *Ant* triple null mice, we independently generated quadruple null mice for *Ant1, Ant2, Ant4* and *Ppif*. We compared the CRC and swelling measurements in liver mitochondria from these quadruple null mice versus *Ant1^-/-^ Ant4^-/-^ Ant2^fl/fl^* control (ANT2 only) mice (Fig. 3A-C). While ANT2 only mitochondria took up a few boluses of 40 μM Ca^2+^ before opening and swelling, the quadruple null never underwent swelling and took up Ca^2+^ boluses until they assay was saturated (Fig. 3A and B). Electron microscopy showed that liver mitochondria from *Ant1^-/-^ Ant4^-/-^ Ant2^fl/fl^* control mice, *Ant1^-/-^ Ant4^-/-^ Ant2^fl/fl-Alb-Cre^* mice, and *Ppif*^-/-^ mice had normal mitochondria at baseline in Ca^2+^ free buffer, but at 800 μM Ca^2+^ all showed profound swelling. Conversely, mitochondria from *Ant1^-/-^ Ant4^-/-^ Ppif^-/-^ Ant2^fl/fl-Alb-Cre^* quadruple null livers failed to show swelling (Fig. 3C) despite our also using half the amount of mitochondria (1 mg versus 2 mg) to correct for baseline absorbance (Fig. S2A). Similar to the *Ant* triple null mitochondria, *Ant1^-/-^ Ant4^-/-^ Ppif^-/-^ Ant2^fl/fl-Alb-Cre^* quadruple null mitochondria displayed no alteration in morphology or stimulated oxygen consumption under energized conditions (Fig. S2B and C). This is the first time that MPT has ever been genetically inhibited, thus demonstrating its molecular existence.

**Fig. 3.**
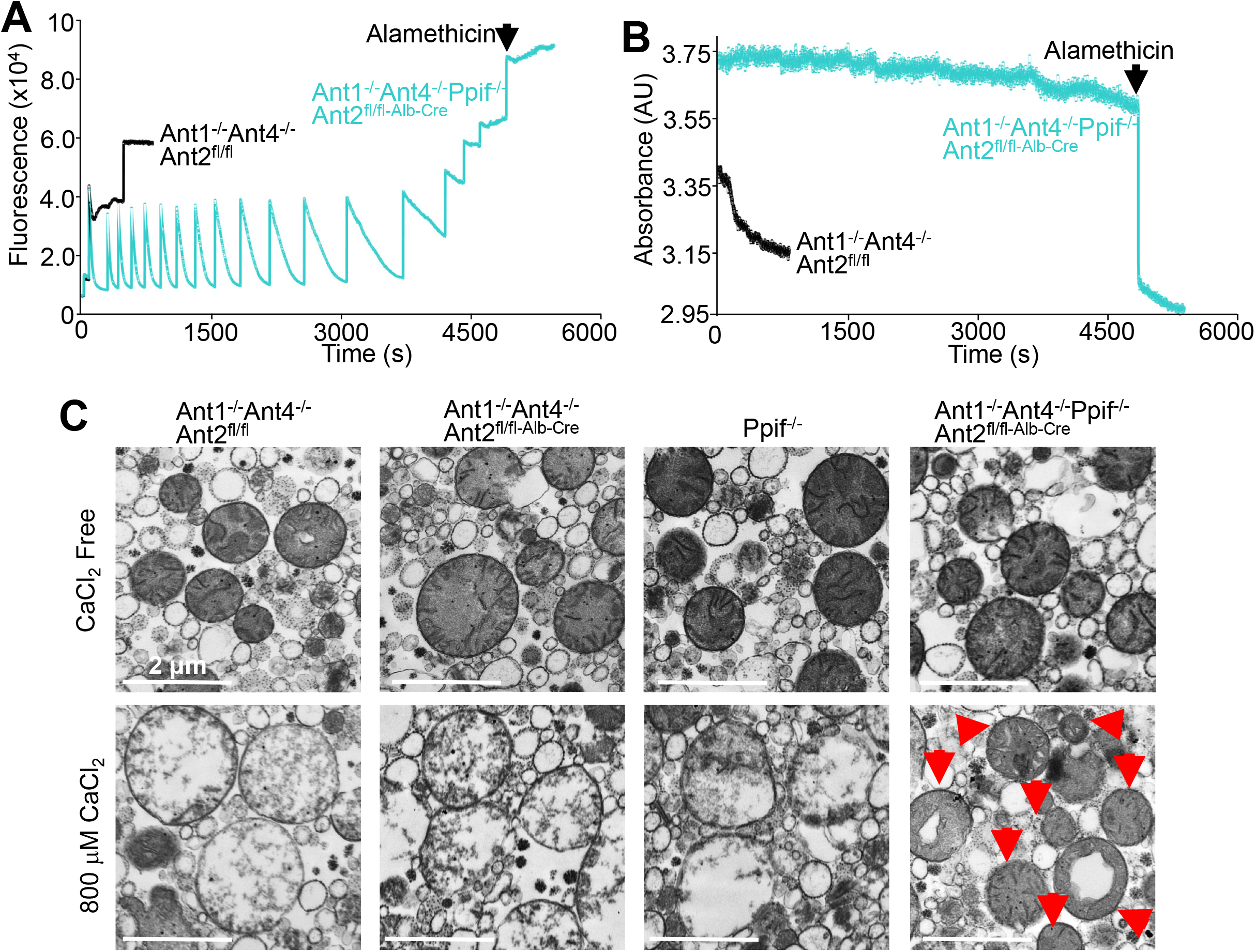
Genetic deletion of the *Ant* family and *Ppif* completely inhibits MPTP opening. (**A**) Ca^2+^ retention capacity assay performed in purified liver mitochondria isolated from *Ant1^-/-^ Ant4^-/-^ Ant2^fl/fl^* (controls that express ANT2 only) or *Ant1^-/-^ Ant4^-/-^ Ppif^-/-^ Ant2^fl/fl-Alb-Cre^* (quadruple KO) mice. Two milligrams of mitochondria per assay were used and 40 μM pulses of CaCl_2_, were continually given until mitochondrial Ca^2+^ uptake no longer occurred or the mitochondria underwent MPT as evidenced by irreversible increase in Calcium Green-5N fluorescence, which was saturated in the presence of alamethicin (arrow head). Similar results were observed across 3 independent assays. (**C**) Electron microscopy images of isolated liver mitochondria treated with or without 800 μM CaCl_2_ from *Ant1^-/-^ Ant4^-/-^ Ant2^fl/fl^* (ANT2 only expressing controls), *Ant1^-/-^ Ant4^-/-^ ANT2^fl/fl-Alb-Cre^* (total *Ant* nulls), *Ppif*^-/-^, and *Ant1^-/-^ Ant4^-/-^ Ppif^-/-^ Ant2^fl/fl-Alb-Cre^* (quadruple null) mice. The red arrows show mitochondria that were refractory to swelling in the quadruple nulls with 800 μM CaCl_2_. Representative images are shown that are consistent across all fields analyzed from 2 independent isolations.

To extend these findings to another model system we also generated mouse embryonic fibroblasts (MEFs) lacking all ANT isoforms. An adenovirus expressing Cre recombinase was used to delete the *Ant2* gene, shown by Western blotting (Fig. 4A). In contrast to the liver mitochondria, mitochondria isolated from *Ant1^-/-^ Ant4^-/-^ Ant2*^*fl/f*-(AdCre)^ MEFs (triple null) did not show compensation in their ability to transport ATP/ADP so that these mitochondria had almost no respiratory capacity (Fig. 4B). However, these MEFs lacking all 3 *Ant* genes still showed mitochondrial membrane potential (ΔΨ) in culture with TMRE fluorescence (Fig. 4C), although this is likely due to support from glycolysis and reverse mitochondrial ATPase activity to maintain ΔΨ. To further explore this concept that *Ant* triple null MEFs depend on glycolysis we grew them in glucose free, galactose-containing culture media making glycolysis an ATP futile process. Under these conditions the *Ant* null MEFs died within 24 hours whereas the control MEFs were unaffected (Fig. 4D).

**Fig. 4.**
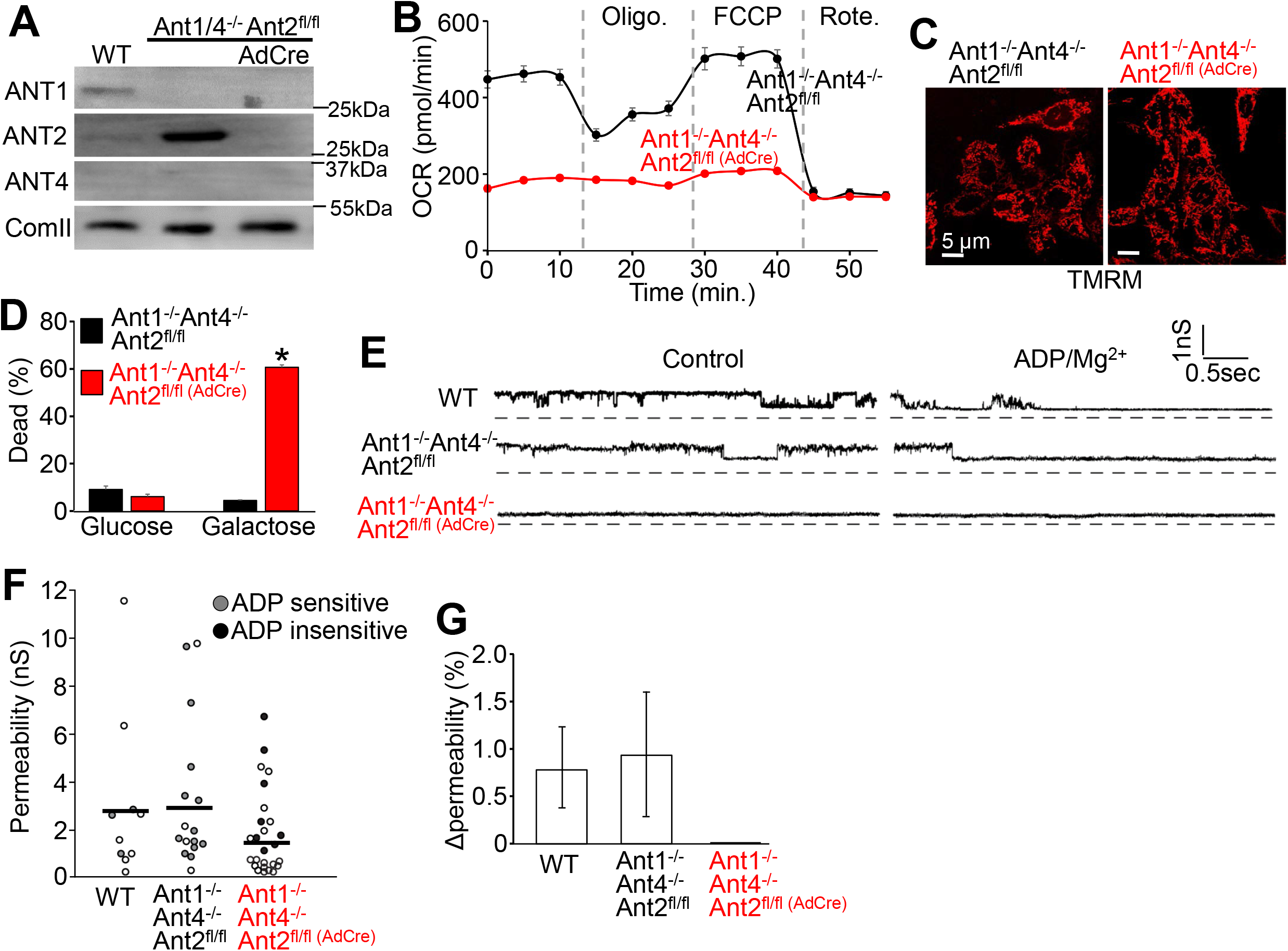
The ANT family is the pore forming component of the MPTP in MEFs. (**A**) Western blots of ANT1, ANT2, ANT4, and loading control (Complex II, ComII) from extracts of isolated mitochondria from MEFs that are wild-type (WT), *Ant1^-/-^ Ant4^-/-^ Ant^fl/fl^*, treated with or without adenovirus expressing Cre recombinase (AdCre will delete *Ant2*). (**B**) Oxygen consumption rate (**OCR**) of stable MEF lines that were *Ant1^-/-^ Ant4^-/-^ Ant2^fl/fl^*, versus *Ant1^-/-^ Ant4^-/-^ Ant2^fl/fl(AdCre)^* (n=5). Respiration was challenged with sequential treatment with 2 μM oligomycin (Oligo), 5 μM FCCP, and 0.5 μM Rotenone (Rote). N=5. (**C**) Confocal microscopic images of TMRE staining of the same MEF lines described in “B”. Similar profiles of TMRE fluorescence were observed in 3 independent experiments. (**D**) Cell death analysis of the *Ant* triple null MEFs versus the *Ant1^-/-^ Ant4*^-/-^ (ANT2 only) controls, subjected to glucose containing (normal) or glucose free, galactose containing media for 24 hours. Cell death was determined by loss of plasma membrane integrity (n=3). (**E**) Representative patch clamp current traces of mitoplasts isolated from the MEFs described in “A” in the presence of 250 μM CaCl_2_. The high conductance openings in the WT and ANT2 only lines were sensitive to addition of 1 mM ADP/Mg^2+^ (right). (**F**) Scatter plot of the permeability in nanosiemens (nS) from all recorded patches. The number of patches analyzed is represented by all the data plots (circles). The bold horizontal lines indicate the average permeability. The solid gray and black circles represent channels that were either sensitive or insensitive to addition of 1 mM ADP/MgCl_2_, respectively. (**G**) Bar histograms of average (± SD) change in permeability after bath perfusion with 1 mM ADP/MgCl_2_.

A final assay that is used to assess MPTP channel activity is patch clamping of the inner mitochondrial membrane to identify the characteristic conductance properties of this inducible pore. Mitoplasts isolated from WT and ANT2 only expressing MEFs (*Ant1^-/-^ Ant4^-/-^ Ant2^fl/fl^*) showed channel opening and conductance levels consistent with the MPTP and importantly ADP, an MPTP suppressor, inhibited all such pore opening events (Fig. 4E and 4F). However, patching of mitoplasts from *Ant* triple null MEFs showed a much lower prevalence of large pores, and all such rare pores were unresponsive to ADP, suggesting that no MPTP like channels were present without ANT (Fig. 4E and G). These results suggest that the ANT family is the primary inner membrane channel forming component of the MPTP in MEFs.

In summary, we have genetically abolished MPT by deletion of the ANT family and CypD in liver mitochondria. This represents the first complete inhibition of this activity and it provides molecular proof that the MPTP exists and is not an artefactual ex vivo phenomenon. Our data support a model of the MPTP consisting of only ANT in MEFs, yet in liver mitochondria it appears to exist as two distinct molecular components, with one being comprised of the ANTs (Fig. S3). In liver mitochondria this ANT containing MPT activity can occur in the absence of CypD, yet a second alternative activity, completely requires CypD isomerase activity. The possibility that the MPTP is composed of at least two pore-forming components may explain why identifying its gene products have been so challenging. In order to identify the CypD-dependent alternative MPTP channel, the ANTs must be accounted for either by genetic deletion or pharmacological inhibition. With this new understanding previously speculated MPTP-forming components, including various other slc25a family members and the mitochondrial F_1_F_O_-ATPase (*18–20, 28, 29*), should be re-evaluated.

## Acknowledgments

We thank Dr Douglas C. Wallace (Children’s Hospital of Philadelphia) for supplying *Ant1*^-/-^ and *Ant2*-loxP targeted mice. We thank Gyorgy Hajnoczky for evaluation of the manuscript.

## Author Contributions

J.K. and J.D.M designed the study and wrote the manuscript. J.K., M.J.B., H.K., M.A.S., N. L., and P.M.P performed experiments and analyzed data. N.T. and D.C.W. shared previously generated mutant mouse lines.

## Data and materials availability

All data are contained within the figures of the paper or supplemental figures, or available upon reasonable request.

## List of supplementary materials

Materials and Methods

Fig S1-S3

References 30,31

