## Supplemental methods and figures for "Inhibition of mitochondrial permeability transition by deletion of the ANT family and CypD"

**SUPPLEMENTAL MATERIALS INCLUDED**

Materials and Methods

Figures S1 - S3

### Materials and Methods

**Animal Models.** *Ant1*$^{-/-}$ *(Slc25a4) Ant2*-LoxP (fl) (*Slc25a5*) mice and *Ant4*$^{-/-}$ (*Slc25a31*) mice were described previously (*16, 17*). To generate liver specific deletion *Ant2*$^{fl/fl}$ in mice we crossed in an albumin-Cre transgenic line (Jackson Laboratory: 003574). To delete the cyclophilin D (CypD) gene product we used mice lacking a functional *Ppif* gene (*27*). *Ppif*$^{-/-}$ mice were crossed to contain the *Ant1*$^{-/-}$ *Ant4*$^{-/-}$ and *Ant2*$^{fl/fl\text{-}Alb\text{-}Cre}$ alleles to generate quadruple gene-deleted mice. Triple *Ant* null liver was generated in *Ant1*$^{-/-}$ *Ant4*$^{-/-}$ *Ant2*$^{fl/fl\text{-}Alb\text{-}Cre}$ mice, which were compared with *Ant1*$^{-/-}$ *Ant4*$^{-/-}$ and *Ant2*$^{fl/fl}$ that lacked the Albumin-Cre transgene, and hence these mice only expressed ANT2 (ANT2 only) which is the only ANT family member normally expressed in liver. However, liver mitochondrial from ANT2 only mice showed identical mitochondrial $Ca^{2+}$ retention capacity and $Ca^{2+}$-induced swelling as fully wild-type (WT) mitochondria (data not shown). Moreover, mitochondrial ultrastructure in liver from *Ppif*$^{-/-}$ mice showed no alterations compared with WT mitochondria, similar to no defect in *Ant1*$^{-/-}$ *Ant4*$^{-/-}$ *Ant2*$^{fl/fl\text{-}Alb\text{-}Cre}$ mice (data not shown, see fig. S2C).

All experimental procedures with animals were approved by the Institutional Animal Care and Use Committee of Cincinnati Children's Medical Center, protocols IACUC 2015-0047 and 2016-0069. We have complied with the relevant ethical considerations for animal usage overseen by this committee. The number of mice used in this study reflects the minimum number needed to achieve statistical significance based on experience and previous power analysis. Blinding was performed for some experimental procedures with mice, although blinding was not possible in every instance. Both sexes of mice were used.

**Cell Culture Models.** Mouse embryonic fibroblasts (MEFs) were generated by harvesting *Ant1*$^{-/-}$ *Ant2*$^{fl/fl}$ *Ant4*$^{-/-}$ embryonic (E) day 10.5 embryos. Following the removal of the internal organs and the head, the remainder of the bodies were passed through a 25 gauge needle and plated on 10 $cm^2$ tissue culture dishes and placed in DMEM (ThermoFisher Scientific) containing 10% Bovine growth serum (BGS, ThermoFisher Scientific), penicillin streptomycin (ThermoFisher Scientific), and non-essential amino acids (Invitrogen). After two passages, the MEFs were then subjected to SV40 large T antigen immortalization by infecting with an SV40 T-antigen expressing adenovirus.

Once immortalization was achieved, cells were treated with an adenovirus expressing Cre recombinase (AdCre) for *Ant2* gene deletion. Before infection these MEFs were switched to “rho zero media” which was the standard DMEM media described above except enhanced with 50 μg/mL uridine (Sigma-Aldrich) and 1 mM sodium pyruvate (ThermoFisher Scientific). Following infection, these MEFs were subjected to FACS to isolate GFP positive cell that was part of the AdCre virus, to establish clonal triple *Ant* deleted MEF lines. Validation of the loss of ANT2 protein was determined by Western blot analysis. To determine if the *Ant* triple null MEFs relied on mitochondrial produced ATP for survival, we cultured the MEFs in glucose free media (ThermoFisher Scientific) containing all the normal supplements with the addition of 4500 mg/ml galactose (Sigma-Aldrich). Twenty-four hours following the replacement of the media we measured cell death using the MUSE Cell Analyzer (MilliporeSigma) and the Muse Count & Viability Assay Kit (MilliporeSigma), which measures membrane permeability to assess dead cells.

**Mitochondrial Isolation.** Brain, heart, muscle, liver, testis and MEF mitochondria were isolated in MS-EGTA buffer (225 mM mannitol, 75 mM sucrose, 5 mM HEPES, and 1 mM EGTA, pH 7.4 (Sigma-Aldrich)). The various tissues were minced into 2 mm X 2 mm pieces and then homogenized used a glass and Teflon tissue homogenizer (8-15 strokes, depending on the tissue) on ice and in MS-EGTA buffer. The MEFs were grown to confluency on 4 Nunc Bioassay Dishes (Sigma-Aldrich) and then harvested and pelleted in MS-EGTA buffer. The cells were then suspended in 7 mL of buffer and homogenized with a glass and Teflon tissue homogenizer (10 strokes). The homogenates were then subjected to a 2,500xg centrifugation for 5 minutes (2X) and the supernatants were then centrifuged at 11,500xg for 10 minutes. Pellets were then washed and centrifuged 2X and then snap frozen or suspended in KCl buffer (125 mM KCl, 20 mM HEPES, 2 mM $MgCl_2$, 2 mM $KH_2PO_4$, and 40 μM EGTA, pH 7.2 (Sigma-Aldrich)).

**Calcium Retention Capacity and Mitochondrial Swelling Assays.** The calcium retention capacity (CRC) assay and the mitochondrial swelling assay were performed simultaneously using a dual-detector (one to measure fluorescence and the other to measure absorbance) single-cuvette based fluorimeteric PTI-system (Horiba Scientific).

Depending on the experiment, 1 mg or 2 mg of isolated mitochondria were loaded into the cuvette along with 250 nM Calcium Green-5N (Invitrogen), 7 mM pyruvate (Sigma-Aldrich), and 1 mM malate (Sigma-Aldrich) and brought up to 1 mL using KCl buffer. Mitochondria were then pulsed with sequential additions of $CaCl_2$ (20, 40, or 400 μM) until MPTP opening occurred or until the mitochondria reached a $CaCl_2$ saturation point and could no longer take up more $Ca^{2+}$. In certain situations where the experimental mitochondria did not undergo a swelling event following $CaCl_2$ additions, they would be subjected to a membrane permeabilizing agent, 40 μM alamethicin (Santa Cruz Biotechnology), to show that they still had swelling capacity and were not otherwise compromised.

**TMRE Assay.** MEFs were plated on glass bottom tissue culture dishes and loaded with 50 nM TMRE (ThermoFisher Scientific) for 10 minutes. Following loading cells were washed twice with Hank's balanced salt solutions (HBSS) (ThermoFisher Scientific) and visualized using a confocal microscopy. Isolated liver mitochondria suspended in KCl buffer were loaded with 50 nM TMRE for 10 minutes then centrifuged and washed with fresh KCl buffer. The mitochondria were then analyzed by flow cytometry. TMRE positivity was confirmed by treating loaded mitochondria with 5 μM FCCP (Abcam) and recording a large negative shift in the entire mitochondrial population.

**Western Blotting.** All Western blots were performed using isolated mitochondrial lysates. Following mitochondrial isolations, the mitochondrial pellets were suspended in radioimmunoprecipitation (RIPA) buffer containing protease inhibitor cocktails (Roche). The samples were then sonicated and the insoluble fractions were discarded following centrifugation. SDS sample buffer was added to the lysates and samples were boiled for 5 minutes. The samples were then loaded onto 10-15% acrylamide gels and then transferred onto PDVF transfer membranes (MilliporeSigma). The following primary antibodies were used: Slc25a4 (ANT1) (SAB; 32484; 1:500), SLC25A5 (ANT2) (Cell Signaling Technology; 14671; 1:500), ANT4 Polyclonal Antibody (SAB; 40596-1; 1:500), and Total OXPHOS Rodent WB Antibody Cocktail (that contained complexes 1-5) (Abcam; ab110413; 1:15,000).

**Metabolic analysis.** Oxidative consumption rate (OCR) was determined using an XF extracellular flux analyzer (Seahorse Bioscience). MEFs were plated to confluency on XF24 cell culture microplates and subjected to the Mito Stress Test Kit (Seahorse Bioscience) using the standard protocol provided. Briefly, after basal respiration was measured the MEFs were treated sequentially with 2 μM oligomycin, 5 μM FCCP, and 0.5 μM rotenone. Isolated mitochondria (300 μg) were loaded onto an XF24 cell culture microplates in KCl buffer containing 1 mM malate and 7 mM pyruvate and subjected to the Mito Stress Test Kit using the standard protocol provided. Briefly, after basal respiration was measured the MEFs were treated sequentially with 10 mM ADP, 2 μM oligomycin, 5 μM FCCP, and 0.5 μM rotenone.

**Histological analyses.** Histological analysis of the hematoxylin and eosin stained slides of 7 μm liver sections were performed using an upright microscope 4X objective. Electron microscopy was performed on livers and isolated mitochondria from livers. Prior to fixation, the isolated mitochondria were subjected to the mitochondrial swelling assay with or without $CaCl_2$ (2 pulses of 400 μM $CaCl_2$ to achieve 800 μM). Samples were then fixed in glutaraldehyde and cacodylate, embedded in epoxy resin, and sectioned. Sections were counterstained with uranyl acetate and lead citrate. Images were taken at 5000X and 3500X.

**Patch clamping techniques.** MEFs at 80% confluency were harvested and mitochondria were isolated as previously described (*30*). Mitoplasts were spontaneously formed by diluting isolated mitochondria 100-fold in patching media (see below) for 10 minutes, which caused controlled swelling and rupture of the outer membrane. Mitoplasts can be easily distinguished from intact mitochondria as they appear as larger and more translucent ring-like structures studded by a curled outer membrane. Inner membrane patches were excised from mitoplasts after formation of a seal using micropipettes with approximately 0.3-μm tips and resistances of 10–30 MΩ at room temperature. Patching media was 150 mM KCl, 5 mM HEPES, 1.25 mM $CaCl_2$, 1 mM EGTA, pH 7.4. Voltages were clamped with a Dagan 1200 amplifier and reported as pipette potentials. Permeability was typically determined from stable current levels and/or total amplitude histograms of 30 s of data at +20 mV. pClamp version 8 (Axon

Instruments) and WinEDR v2.3.3 (Strathclyde Electrophysiological Software) were used for current analysis as previously described (31). Sample rate was 5 kHz with 1–2 kHz filtration. To determine if the large conductance activity was due to the mitochondrial permeability transition pore through the ANT family patching media containing 1 mM ADP/$MgCl_2$ was administered by bath perfusion.

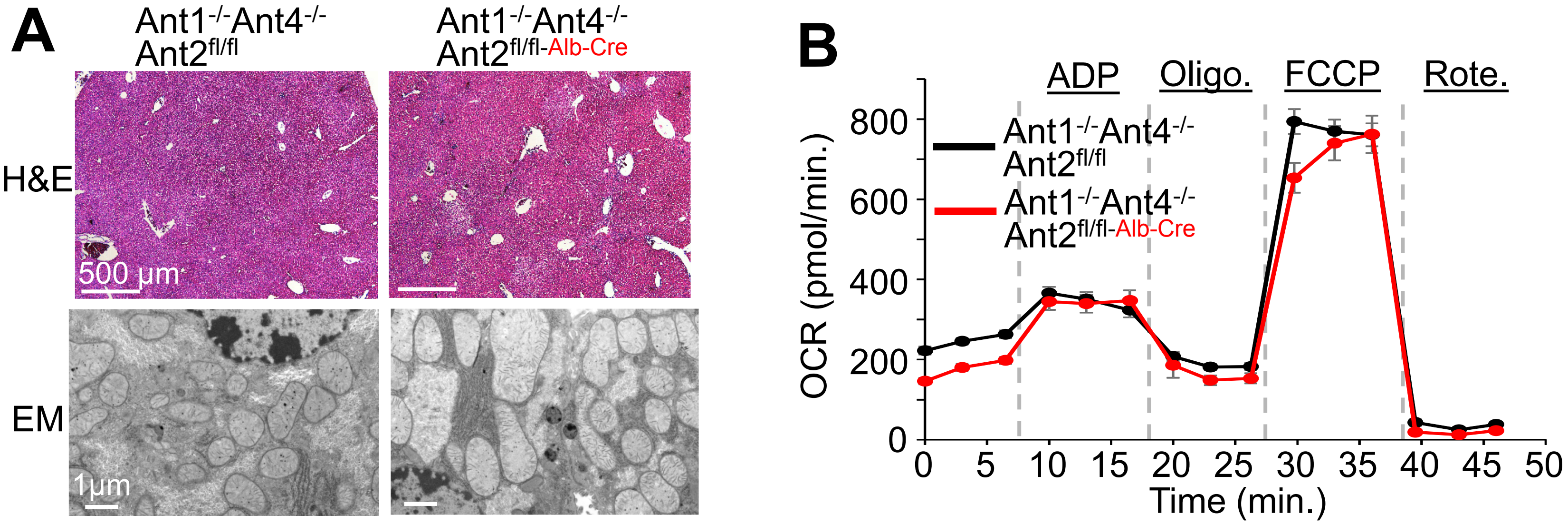


**Fig. S1. Loss of *Ant1/2/4* genes in the liver does not compromise mitochondrial energetics or cause tissue pathology**. (**A**) Representative histological images of H&E-stained liver sections from *Ant1*$^{-/-}$ *Ant4*$^{-/-}$ *Ant2*$^{fl/fl}$, versus *Ant1*$^{-/-}$ *Ant4*$^{-/-}$ *Ant2*$^{fl/fl\text{-}Alb\text{-}Cre}$ mice (Top) and representative electron microscopy images of liver sections from the same mice. (**B**) Oxygen consumption rate (OCR) of isolated liver mitochondria from mice described in "A" (n=4). Mitochondria were treated sequentially treated with 10 mM ADP, 2 µM oligomycin (Oligo.), 5 µM FCCP, and 0.5 µM rotenone (Rote.).

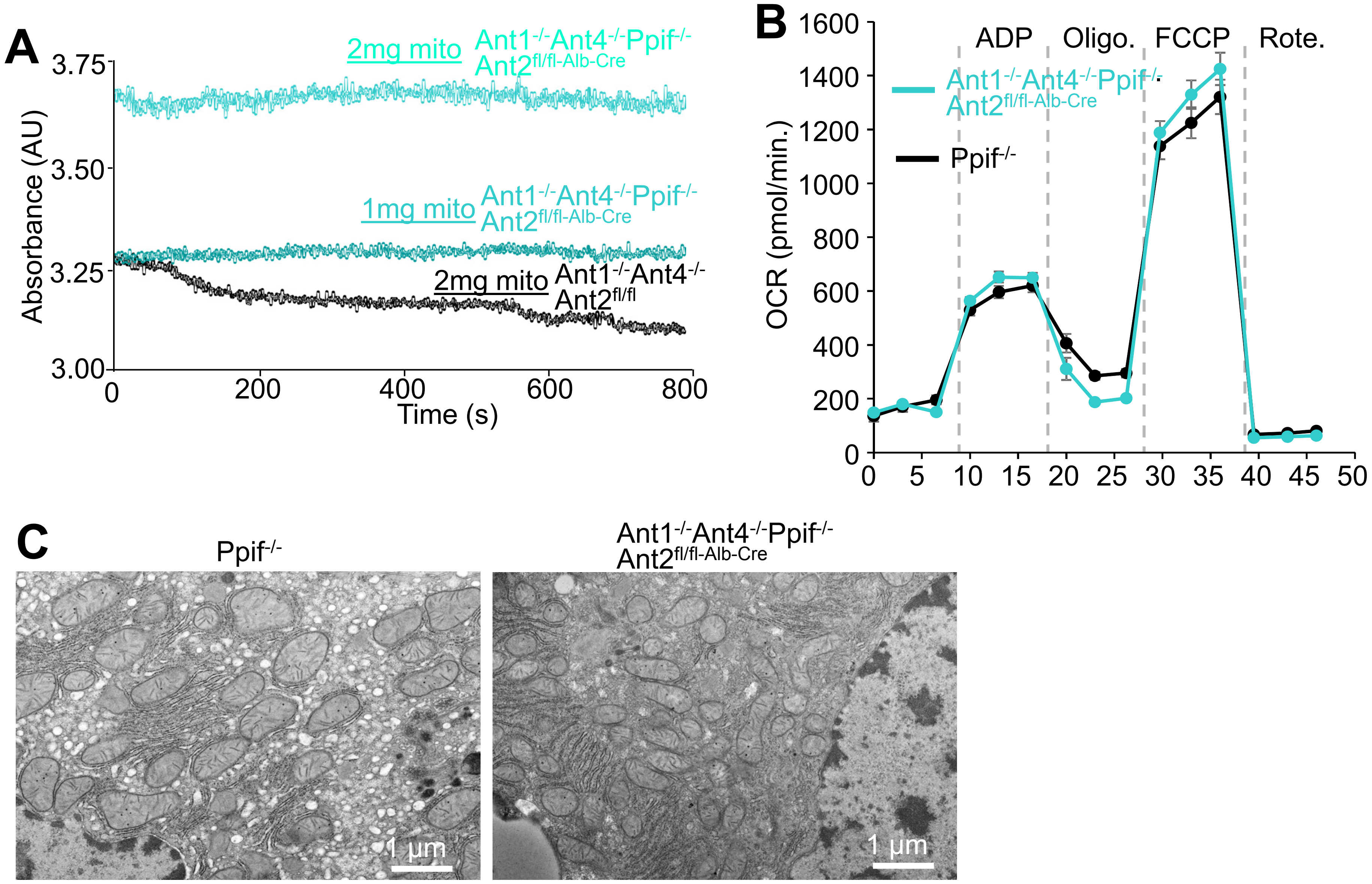
A
Absorbance (AU)
3.75
3.50
3.25
3.00
0
200
400
600
800
Time (s)
2mg mito Ant1-/-Ant4-/-Ppif-/-Ant2fl/fl-Alb-Cre
1mg mito Ant1-/-Ant4-/-Ppif-/-Ant2fl/fl-Alb-Cre
2mg mito Ant1-/-Ant4-/-Ant2fl/fl
B
OCR (pmol/min.)
1600
1400
1200
1000
800
600
400
200
0
0
5
10
15
20
25
30
35
40
45
50
ADP
Oligo.
FCCP
Rote.
Ant1-/-Ant4-/-Ppif-/-Ant2fl/fl-Alb-Cre
Ppif-/-
C
Ppif-/-
Ant1-/-Ant4-/-Ppif-/-Ant2fl/fl-Alb-Cre
1 µm
1 µm

**Fig. S2. Mitochondria from quadruple null mice fail to undergo $Ca^{2+}$ induced MPTP, although no defects in energetics or mitochondrial ultrastructure were observed**. (**A**) Mitochondrial swelling measured by light absorbance performed on liver mitochondria isolated from *Ant1*$^{-/-}$ *Ant4*$^{-/-}$ *Ant2*$^{fl/fl}$ (ANT2 only expressing controls) or *Ant1*$^{-/-}$ *Ant4*$^{-/-}$ *Ppif*$^{-/-}$ *Ant2*$^{fl/fl\text{-}Alb\text{-}Cre}$ (quadruple KO) mice. Swelling was initiated with 3 pulses of 400 µM $CaCl_2$. Two milligrams of ANT2 only control mitochondria showed swelling with this very robust $Ca^{2+}$ challenge but neither 2 mg nor 1 mg of quadruple KO mitochondria swelled, regardless of baseline starting absorbance. (**B**). Oxygen consumption rate (OCR) of isolated liver mitochondria from *Ant1*$^{-/-}$ *Ant4*$^{-/-}$ *Ant2*$^{fl/fl}$ (ANT2 only expressing controls) or *Ant1*$^{-/-}$ *Ant4*$^{-/-}$ *Ppif*$^{-/-}$ *Ant2*$^{fl/fl\text{-}Alb\text{-}Cre}$ (quadruple KO) mice (n=4). Mitochondria were treated sequentially treated with 10 mM ADP, 2 µM oligomycin (Oligo.), 5 µM FCCP, and 0.5 µM rotenone (Rote.). (**C**) Representative transmission electron microscopy images of liver histological sections from Ppif$^{-/-}$ mice versus *Ant1*$^{-/-}$ *Ant4*$^{-/-}$ *Ppif*$^{-/-}$ *Ant2*$^{fl/fl\text{-}Alb\text{-}Cre}$ (quadruple KO) mice. No ultrastructural abnormalities were observed in liver mitochondria from either genotype. All representative data shown in this figure are from at least 3 independent assays.

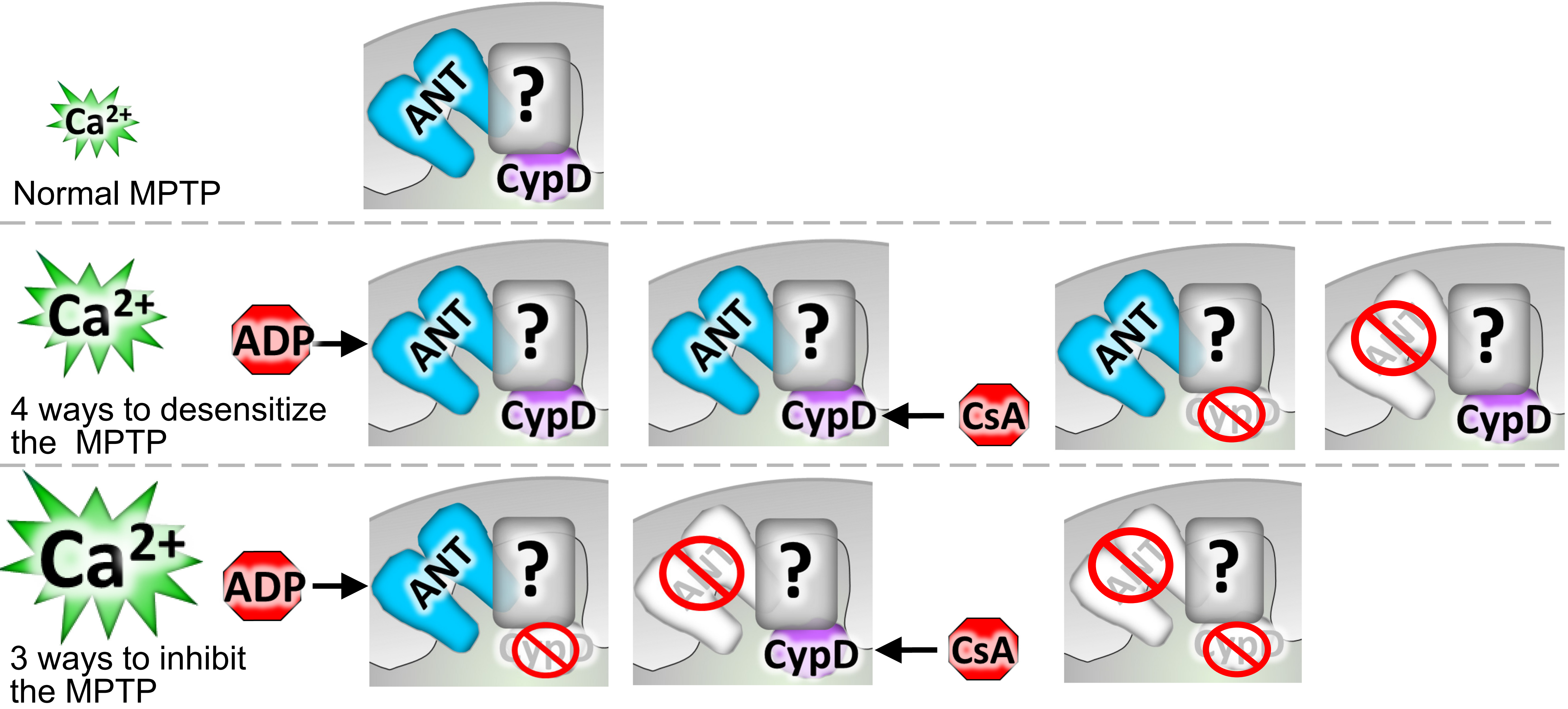


**Fig. S3 Model of the MPTP.** Schematic representation of the mitochondrial permeability transition pore consisting of the ANT family, CypD, and an additional unknown inner membrane pore forming component (?). Normally the MPTP opens in the presence of $Ca^{2+}$ stimulus. Upon genetic removal of the ANT family or CypD (*Ppif* gene) or pharmacological inhibition of the MPTP through ADP or CsA, the MPTP becomes desensitized to $Ca^{2+}$-induced opening, but with high enough levels of $Ca^{2+}$ MPTP opening can still occur. However, pharmacological inhibition of the MPTP with ADP or CsA combined with genetic removal of CypD or the ANT family greatly desensitizes or completely inhibits the MPTP to $Ca^{2+}$-induced opening. Genetic removal of both the entire ANT family and CypD completely inhibits $Ca^{2+}$-induced MPTP opening.
